# Microbial activated mineral weathering and cementation as precursors to hardpan formation and heavy metal encapsulation in sulfidic tailings

**DOI:** 10.1101/2020.09.07.285858

**Authors:** Yunjia Liu, Songlin Wu, Gordon Southam, Ting-Shan Chan, Ying-Rui Lu, David J. Paterson, Longbin Huang

## Abstract

Extensive mineral weathering and formation of large amounts of Fe-rich secondary mineral gels have been identified as precursors critical to forming massive hardpan caps in the surface layers of sulfidic tailings. However, how to initiate and accelerate these precursor processes remains to be established before developing this hardpan-based novel method to rehabilitate sulfidic tailings landscapes. In a 5-month microcosm experiment, the present study has demonstrated the concept of bio-engineering sulfidic tailings by inoculating Fe/S-oxidizing bacterial consortium to accelerate the weathering of sulfides and other Si-rich minerals for mineral gels formation. Synchrotron-based X-ray absorption fine structure spectroscopy (XAFS) demonstrated that the weathering of pyrite and biotite-like minerals was rapidly accelerated by the presence of Fe/S-oxidizing bacterial consortium. The microbial process and associated mineral transformation led to the formation of critical precursor mineral gels, *i*.*e*., jarosite-like minerals, as indicators of the onset of hardpan formation. In the meantime, the labile Zn liberated in the weathering was encapsulated in the jarosite-like minerals as revealed by X-ray fluorescence microscopy (XFM). This concept-proven bio-engineering process is ready to be scaled up in further studies under field conditions to develop an alternative hardpan-based method to cover and rehabilitate sulfidic tailing landscapes.

**TOC Art:** 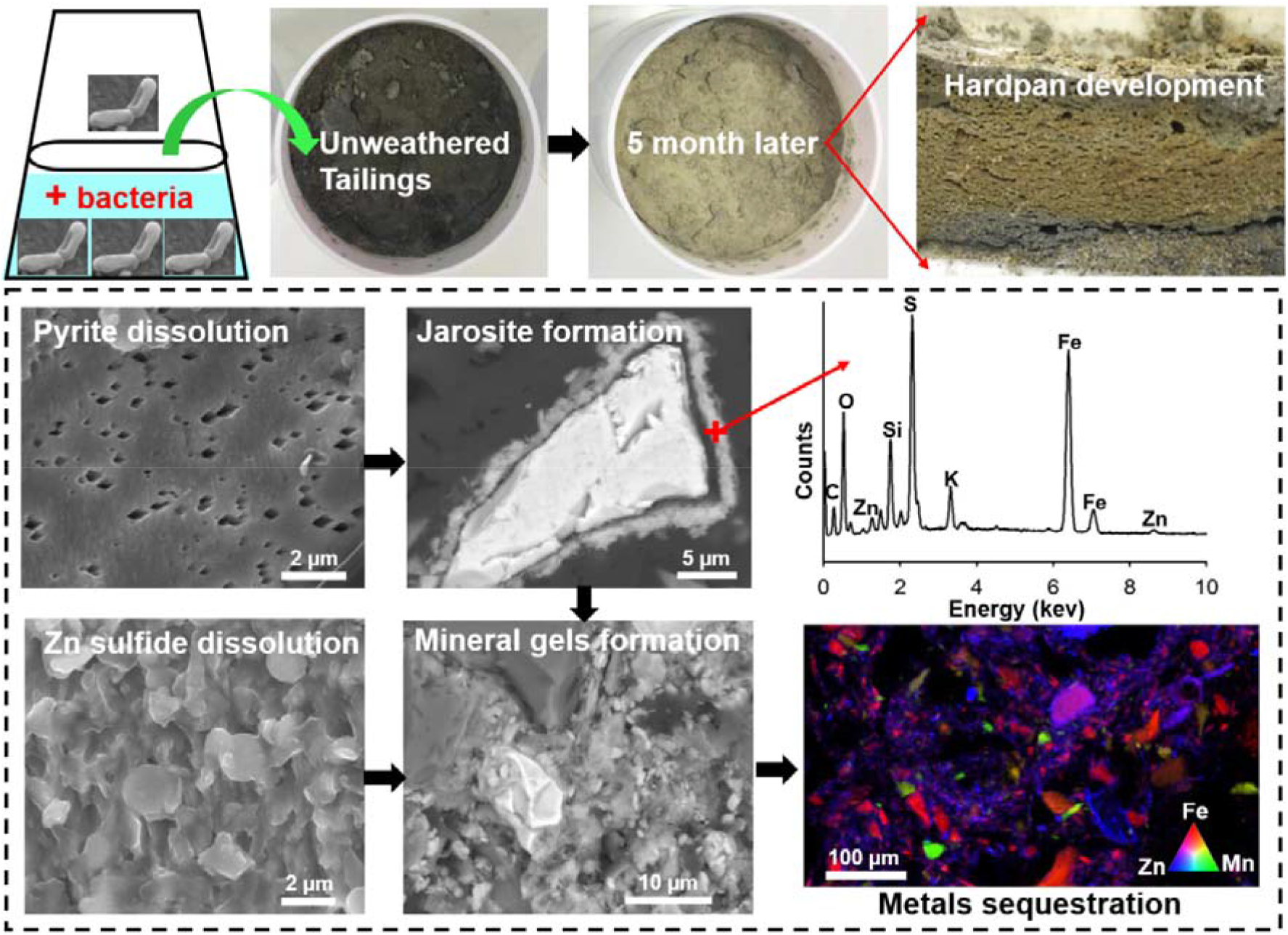

## 1 Introduction

Sulfidic and metallic tailings contain abundant reactive minerals such as pyrite (FeS_2_), pyrrhotite (Fe_1-x_S), sphalerite (Zn[Fe]S) and galena (PbS), posing high risks of metallic drainage pollution^1, 2^. Successful closure of sulfidic tailings storage facilities (TSF) requires not only physical isolation of the tailings from water and oxidative conditions, but also hydrogeochemical stabilization in the mineral phase to avoid secondary risks of metal(loid) dissolution^3^. So far, rehabilitation and closure of sulfidic TSF have commonly adopted conventional cover systems with installed capillary break layers. These caps are installed with compressed clay and bengin rocks, which are expensive and limited by material deficiencies and engineering variability under challenging climatic conditions^4^. Our recent investigation of a long-term field trial (>15 years) discovered a thick and continuous layer of hardpan cap (30 – >50 cm thick) naturally formed at the surface of sulfidic Cu-Pb-Zn tailings^5^. The hardpan cap, together with the cover soil formed hardpan-based duplex soil systems supporting native vegetation under semi-arid climatic conditions^5, 6^. The massive hardpan layer provided physical structure of capillary break between root zones above and the reactive tailings underneath^4^. The hardpan also exhibited a high degree of hydrogeochemical stability, since sulfidic minerals herein were extensively weathered and heavy metals were encapsulated in the cemented structure^7^. As a result, a novel cover method, namely hardpan-based duplex soil system has been proposed to overcome the limitations in conventional cover method.

However, natural formation of hardpans may take several decades for achieveing structural ingetrity, mineralogical polymerization and geochemical inertness^4^. Whether this novel concept could be translated into future cover method for rehabilitating sulfidic tailings would depend on if we could establish the engineering process to initiate and accelerate hardpan formation in unweathered sulfidic tailings. The weathering of sulfides and Si-rich minerals to form large amounts of secondary mineral gels is one of the fundamental processes to cement tailings particles, which are in fact a typical geological indicator of hardpan formation^8, 9^. Besides, the secondary mineral gels within the Fe-rich cement layers in the hardpan profile exhibit strong capacities for sequestration of heavy metals (such as Pb and Zn) in sulfidic tailings^5, 7, 10-12^.

Tolerant extremophiles (such as the genus *Acidithiobacillus*: *A. thiooxidans* and *A. ferrooxidans*) which are indigenous to sulfidic tailings, can catalyse the oxidation of Fe^2+^ and S^2-^ in pyrites and other metal sulfides at a much higher rate than chemical oxidation^13-15^. The rates of sulfur oxidation by *A. ferrooxidans* are relatively slow, compared to *A. thiooxidans* at acid pH conditions^16^, but the combined functions of those two microbial species would significantly enhance the oxidation rates and continual oxidation of sulfide minerals by restricting the formation of intermediate precipitates of S^0 17^. The microbes initiated weathering process generates large amounts of secondary minerals^18^, even under the initial circumneutral pH conditions^19, 20^. Several studies have demonstrated that the formation of secondary Fe-bearing minerals driven by *Acidithiobacillus spp*., such as jarosite, schwertmannite, and ferrihydrite^21-24^. As a result, it is proposed here to establish if the Fe/S-oxidizing microbes could be activated to accelerate the weathering processes for secondary mineral gels formation and hardpan development in the sulfidic tailings.

Therefore, the present study has aimed to establish the concept-proof of the bio-engineering process, by harnessing microbial power to accelerate mineral weathering, secondary mineral gels development, as well as the heavy metal stabilization in freshly deposited sulfidic tailings. It was hypothesized that Fe and S oxidizers (a mixure of *A. ferrooxidans* and *A. thiooxidans*) would accelerate the weathering of primary Fe-bearing minerals (*e*.*g*., pyrite and biotite) and formation of secondary mineral gels (*e*.*g*., secondary Fe (oxyhydr)oxides or hydroxysulfate) for cementing mineral particles in the tailings. It is expected that the biotite-like minerals would release interlayer K^+^ to be concurrently incorporated as a structural cation in jarosite under the bioweathering process. These secondary mineral gels would sequester the labile metals released from the weathering due to their strong metal sorption capacity. A suit of microspectroscopic technologies including synchrotron-based X-ray absorption fine structure spectroscopy (XAFS) and X-ray fluorescence microscopy (XFM) coupled with X-ray absorption near edge fine structure spectroscopy (XANES) were employed to examine mineralogical/microstructure changes, secondary Fe-mineral formation and heavy metal sequestration in the neoformed hardpans.

## 2 Materials and methods

### 2.1 Materials

The sulfidic tailings were bulk-sampled at an operating tailings storage facility (TSF) of Cannington mine in Northwest Queensland, Australia (21°85’S, 140°90’E). Tailings geochemistry background information are provided in the **Supporting Information** (**SI**) (**Table S1**). The bulk tailings were oven-dried at 40°C, ground manually and sieved through a 1 mm stainless steel sieve. Approximately 120 g tailings were loaded into an autoclaved Buchner Funnel (70 mm in diameter, 40 mm in height), which was bottom-lined with a 60-µm pore-size polyamide mesh for preventing the loss of the tailing particles. The bacterial consortium used to inoculate the tailings contained *A. ferrooxidans* (DSM 14882) and *A. thiooxidans* (ATCC 19377), which were added into each of the columns at the rate of approximately 2 × 10^7^ cells g^−1^ tailings for *A. ferrooxidans* and 1 × 10^7^ cells g^−1^ tailings for *A. thiooxidans*. The inoculum was suspended in 9k medium^25^ at pH□2.3, containing 0.4 g L^−1^ (NH_4_)_2_SO_4_, 0.1□g L^−1^ K_2_HPO_4_ and 0.4□ g L^−1^ MgSO_4_·7H_2_O, at the rate of 30□ml solution per funnel for each wet-dry cycle. A set of sodium lamps (80 W, 12h light/dark cycling) were installed above the funnel columns to stimulate evaporative processes under field conditions (see **Fig. S1**, experimental setup).

### 2.2 Experimental design

The microcosm study included three treatment factors (with 4 replicates per treatment): the tailings inoculated by the bacterial consortium (containing both *A. thiooxidans* and *A. ferrooxidans*) in the 9k medium (“TB”), the tailings receiving the same medium without bacterial inoculation as the control (“TC”), and time “0” tailings (“T0”, original tailings without any inputs), respectively. Each treatment had four replicates. For the abiotic control, the 9k medium contained 0.1 g L^−1^ sodium azide to suppress bacteria growth. *Acidithiobacillus thiooxidans* and *A. ferrooxidans* in the mid-log growth phase were separately cultured for 14 and 5 days in the 9k medium solution. The cells in the enriched suspension were first separated from the S^0^ and Fe precipitates by filtering the culture solution through a sterile filter paper (Whatman® Grade 1). The cell concentrates in the inoculum were assessed by using a Petroff-Hauser counting chamber (Hausser Scientific, Horsham, PA, USA) and phase-contrast microscope (Nikon Instruments, Tokyo, Japan). Cell viability of *A. thiooxidans* and *A. ferrooxidans* in the inoculum stocks was 85% and 90% respectively, as determined by a Bac-light® fluorescence microscopy (ThermoFisher, Waltham, MA, USA). During the experimental period, an aliquot of 30 ml 9k medium was added to each column every 1-2 days, which maintained the moisture level in a range from 6% to 30% (< the maximal water holding capacity, ca. 32%) in the each wet-dry cycle. The leachate solutions were collected monthly by adding 35 ml 9k medium to each column to collect approximately 2-3 ml for pH, EC, and elemental analysis. Based on monitoring leachate geochemistry, the experiment ended in 5 months after commencement. The tailings from the columns were sampled for physicochemical analysis, from which subsamples were freeze-dried and ground for and micro-spectroscopic analysis such as XRD and synchrotron based XAFS analysis.

### 2.3 Physicochemical analysis

#### Leachate analysis

The pH and EC in the leachate were monitored using a micro pH probe and a conductivity flow-cell (Horiba Ltd. Kyoto, Japan), respectively. Elemental concentrations in the leachates were filtered through 0.45 μm single-use syringe filters and measured using inductively coupled plasma-optical emission spectrometry (ICP-OES, 720ES, Varian, Palo Alto, CA, USA).

#### Chemical sequential extraction

A seven-step sequential extraction method was adopted to examine Zn speciation^26, 27^. The fractions include: (1) water-soluble fraction; (2) exchangeable fraction (targeting adsorbed and exchangeable ions); (3) weakly acid-soluble fraction (targeting carbonates); (4) weakly reducible fraction (targeting amorphous or poorly crystalline (oxyhydr)oxides of Fe^3+^ and Mn^4+^); (5) strongly reducible fraction (targeting crystalline Fe^3+^ and Al^3+^ (oxyhydr)oxides); (6) oxidizable fraction (targeting primary metal sulfides) and (7) residual fraction. Briefly, aliquots of 1.0 g of the air-dried tailing (sieved through 1 mm mesh sieve) were added to 50 mL polypropylene centrifuge tubes. Designated extracting solution was added into the tubes and mixed thoroughly by means of an end-over-end shaker. After each extraction step, sample solutions were centrifuged and the supernatant was further filtered (0.45 μm filtered) into 10 mL polyethylene vials. The solid residue was subsequently rinsed between each extraction step by adding 20 mL of deionised water (Milli-Q, 18.2 MΩ.cm at 25 °C ultrapure water). The filtered solutions were analysed for elements by means of ICP-OES. The detailed extraction steps, conditions and chemicals are listed in **SI** (**Table S2**).

### 2.4 Micro-spectroscopic analysis

#### XRD analysis

The freeze-dried tailing samples were ground into fine powder (< 30 µm) for XRD analysis. XRD traces were obtained from 2θ=10° to 70° and Cu Kα radiation at 40 kV and 30 Ma (Bruker AXS D8, Karlsruhe, Germany). The step size was 2θ=0.02° with a 2 second per step count time. Minerals were identified from the XRD traces using SIeve+ 2018 software and the International Centre for Diffraction Data’s PDF-4/Minerals database.

#### Fe K edge XAFS analysis

The freeze-dried tailing samples were ground into fine powder (< 30 µm) for synchrotron-based Fe K edge (7,112 eV) XAFS analysis. The spectra were collected on beamline 01C1 at the National Synchrotron Radiation Research Centre (NSRRC) in Hsinchu City, Taiwan. Twenty different Fe compounds were used as reference standards (**Fig. S2**). The Fe K-edge XAFS spectra were analysed using ATHENA from the IFEFFIT software package^28^. Detailed methods are described in **SI**. Both Fe K edge extended X-ray absorption fine structure (EXAFS) and X-ray absorption near edge structure (XANES) employed to predict the Fe speciation by using linear combination fitting (LCF). It is essential to note that the sum of components was not set to 100% in the LCF-EXAFS analysis, as there might be other Fe species present that not identified in the fitting. The quality control of the fitting was confirmed by minimal R-factor, Chi-square and Reduced Chi-square.

#### BSE-SEM-EDS examination

The air-dried tailing samples were embedded into the resin and polished for examination of morphology and element composition by using scanning electron microscope (SEM) coupled with energy dispersive spectroscopy (EDS). The unpolished air-dried samples were mounted onto separate 12 mm aluminium stubs with carbon adhesive tabs. Both polished and comparable unpolished samples were coated with approximately 10-20 nm carbon, using a carbon coater (Quorum Q150T, Sussex, UK). The samples were then examined using a SEM (JEOL JSM 6610, Tokyo, Japan) for secondary electron (SE) and back-scattered electron (BSE) imaging. The EDS spectra were collected at 10-15 kV. Aztec energy-acquisition software (Oxford Instruments Version 3.2, Abingdon, UK) was used to analyse the EDS data.

#### XFM and fluorescence-XANES imaging

The air-dried tailing samples were polished and mounted on quartz slides for synchrotron-based XFM analyses at the XFM beamline at the Australian Synchrotron (Melbourne, Australia). The petrographic thin sections (30 µm thickness) were attached to a Perspex sample holder using Mylar tape and mounted on the translation stages of the X-ray microprobe. Incident X-rays, at an energy of 18.5 keV, were selected using a Si (111) monochromatic X-ray beam with Kirkpatrick-Baez mirrors focussing the beam to approximately 2 µm × 2 µm^29^. The X-ray fluorescence emitted by the specimens was collected using the 384-element Maia detector located in a backscatter geometry^30^. The total photon flux was approximately 5.3 ×10^8^ photon per second. The thin sections were raster scanned, moving continuously in the horizontal direction (“on the fly”). The fluorescence-XANES image consisted of a stack of 125 individual maps that were collected at decreasing incident energies from 9.804 to 9.63 keV, across the Zn K-edge (9.659 keV). Within this energy range, the following energy decrements were utilized: (i) 10 eV from 9.804 to 9.724 keV (nine energies), (ii) 2 eV from 9.724 to 9.7 keV (12 energies), (iii) 0.5 eV from 9.7 to 9.65 keV (100 energies), (iv) 5 eV from 9.65 to 9.63 keV (four energies). For these XANES stacks, each of the 125 individual maps was collected using a sampling interval of 2 µm and a dwell time of 2 ms per pixel. The raw data of elemental distribution were analysed using GeoPIXE and the images were generated using the Dynamic Analysis method^31^. The Zn fluorescence-XANES stacks were analysed by the GeoPIXE ‘energy association’ module to identify and select areas where the speciation varied by comparing the intensity ratios between two energies for each pixel. For these various pixel populations, the XANES spectra were extracted from the XANES stack. Thirteen different Zn compounds were used as reference standards (**Fig. S3**). LCF was used to predict the Zn speciation as described above.

### 2.5 Statistical analysis

The physicochemical characteristics were analysed for their significant differences among different treatments by one-way ANOVA at p <0.05 (SPSS Version 20, IBM, Armonk, USA).

## 3 Results

### 3.1 Leachate geochemistry in response to microbial inoculation

During the 5 months of column incubation period, the pH in the tailings remained relatively stable at 2.5 to 3.0 at each sampling cycle (**Fig. S4**), but the leachate EC in both TB and TC increased from about 10 mS cm^−1^ to 20 mS cm^−1^ after the first month of incubation (**Fig. S5**). In the first three months, K concentrations in the leachates increased from 50 mg L^−1^ to 900 mg L^−1^ in the TB, but then decreased rapidly to 33 mg L^−1^ at the end (**Fig. S6**). Concentrations of other key elements (including Si, Zn, Co, Mn, Cd, Fe, and Pb) in the leachate of the TB treatment significantly increased by the third month, but not the TC. Particularly, Zn concentrations in the TB leachates increased to > 1000 mg L^−1^ at the end of the experiment. In contrast, Zn concentrations in the TC leachates remained at 1-20 mg L^−1^ throughout the experiment.

### 3.2 Transformation of Fe-bearing minerals revealed by XRD

The mineralogy of the Fe-bearing phases was significantly altered by the colonization of Fe/S oxidizing bacterial consortium. According to the XRD spectra, jarosite (at 2θ=14.8°, 17.3°, 24.2° and 28.9°, respectively) was only detected in the TB, but not in the TC and T0 (**Fig. 1**). Correspondingly, the intensity of pyrite (at 2θ=33.0°, 37.0° and 40.7°, respectively) in the TB was lower than those in the TC and T0. However, no significant differences were found in Si-rich minerals and carbonates in the tailings among the three treatments. As shown in the XRD analysis, the overall mineral composition of the tailings was composed of Si-rich minerals (*e*.*g*., quartz), sulfides (*e*.*g*., pyrite and sphalerite), and carbonate minerals (calcite and siderite).

**Fig. 1.**
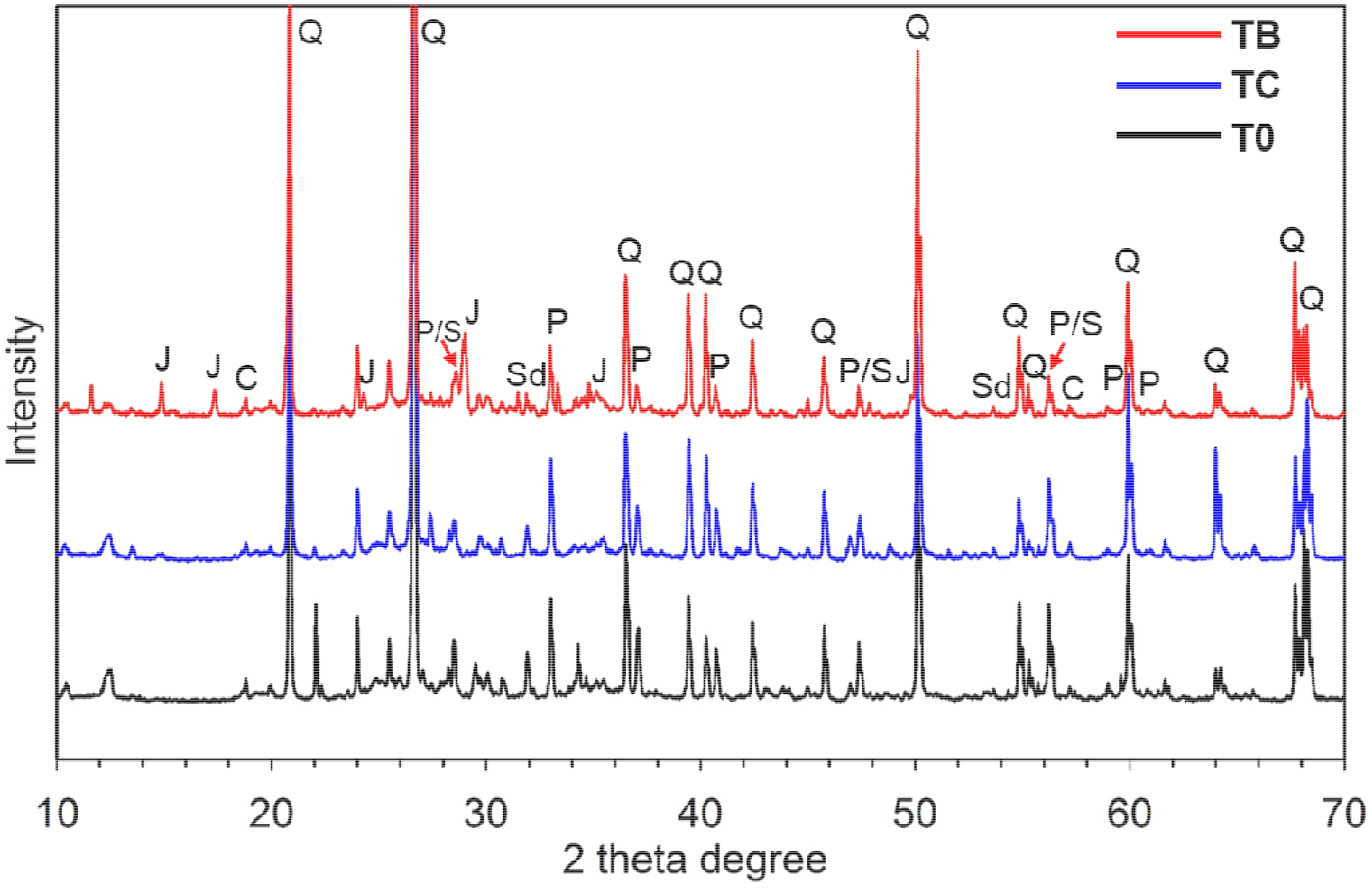
XRD spectra of the tailings. Note: “TB” represents tailings inoculated by the bacterial consortium (containing both *A. thiooxidans* and *A. ferrooxidans*); “TC” represents the tailings receiving the same medium without bacterial inoculation as the control; “T0” represents the time “0” tailings (original tailings). Note: J: Jarosite (KFe_3_(SO_4_)_2_(OH)_6_); C: Calcite (CaCO_3_); Q: Quartz (SiO_2_); P: Pyrite (FeS_2_); S: Sphalerite (Zn[Fe]S); Sd: Siderite (FeCO_3_).

### 3.3 Transformation of Fe phases revealed by XAS analysis

In the pre-edge of Fe K edge XANES, Fe(III) (*e*.*g*., ferrihydrite) was characterized by adsorption peaks around 7113.5 eV, while Fe(II) (*e*.*g*., pyrite) was characterized around 7112.2 eV and 7120.5 eV (**Fig. 2A**). The samples of TC and T0 treatments had pre-peaks at 7112.0 eV in the first derivative of the normalized adsorption, indicating a major occurrence of Fe(II). In contrast, the shoulder of the first derivative of the normalized adsorption was slightly shifted to 7113.5 eV in the TB, indicating more Fe(III) was present in the TB than that in TC and T0. The *k* space of Fe K edge EXAFS spectra demonstrated distinct peak oscillations and line shapes among the three treatments (**Fig. 2B**). The LCF-EXAFS analysis revealed that pyrite and biotite contents were deceased, while the jarosite-like minerals were increased in the TB (**Fig. 2C**). In detail, Fe was present mainly as jarosite (26.9%), ferrihydrite (25.3%), biotite (22.7%), and pyrite (18.8%) in the TB. In contrast, no jarosite was detected in the TC and T0 except for low amounts of ferrihydrite (9.6%) was detected in the TC. The pyrite, biotite, and siderite remained as the dominiant Fe-bearing minerals in the TC and T0, which accounted for more than 80% of the total Fe species. Similarly, the Fe K-edge XANES spectra and LCF-XANES estimated that the increased propotions of jarosite and ferrihydrite in the TB were consistent with the decreases of pyrite and biotite proportions, compared with those in TC and T0 (**Fig. S7**).

**Fig. 2.**
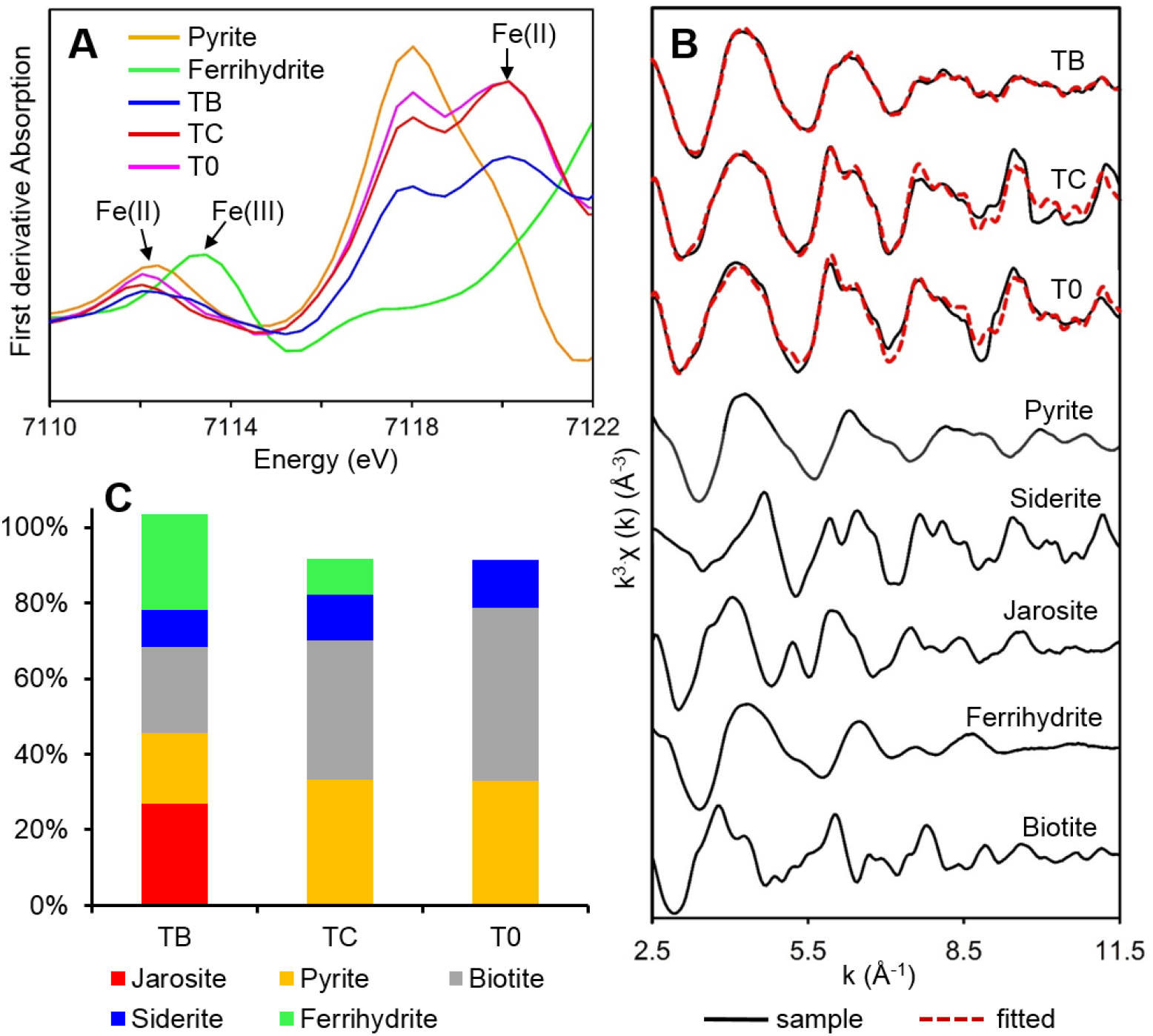
First derivative of absorption of pre-edge of Fe K edge XANES (**A**). *k* space Fe K edge EXAFS spectra showing the distinct Fe phases within the tailings (**B**). Reference Fe minerals are also included in the spectra. LCF-EXAFS prediction of the composition of Fe phases (**C**). Note: “TB”, “TC”, and “T0” are treatments labelled in **Fig 1**. Note: the sum of components was not 100% in the LCF-EXAFS. The r-factor, chi-square and reduced chi-square were 0.0079229, 0.37174, and 0.0021122 for TB; 0.0765859, 2.93539, and 0.0165841 for TC; 0.0732569, 2.96860, and 0.0166775 for T0, respectively.

### 3.4 Bio-weathering induced morphological changes of Fe-bearing minerals

The surface morphology of pyrite particles in the tailings was significantly modified by colonizing bacterial cells. BSE-SEM imaging and EDS spectra demonstrated that massive corrosion pits and precipitates were present at the surfaces of pyrite grains **(Fig. 3A&B)**. The EDS spot analysis demonstrated that elemental composition of the precipitates included Fe, O, C, Al, Si, P, S and K. The magnification of surface area revealed that the size of each corrosion pit was approximately 0.5 µm (**Fig. 3C**). In contrast, the pyrite surface morphology remained largely unchanged with little traces of mineral dissolution or precipitation in the TC **(Fig. S8)**. According to the BSE-SEM examination of polished TB tailings, the pyrite grain was encapsulated within an approximately 1-2 µm thick layer of jarosite-like minerals, forming a mineral passivation layer (**Fig. 3 D**) and cementing tailing particles together (**Fig. 3 E&F**). The examination of unpolished tailings indicated that the cement clusters (each particle is *ca*. 1-2 µm) had the tendency to form microaggregates (*ca*. 100-200 µm) in the TB (**Fig. 3 G-I**). Moreover, the EDS spot analysis revealed that the jarosite-like cements may have embedded metal cations in the TB, such as the labile Zn released from the sulfides undergoing bio-weathering processes.

**Fig. 3.**
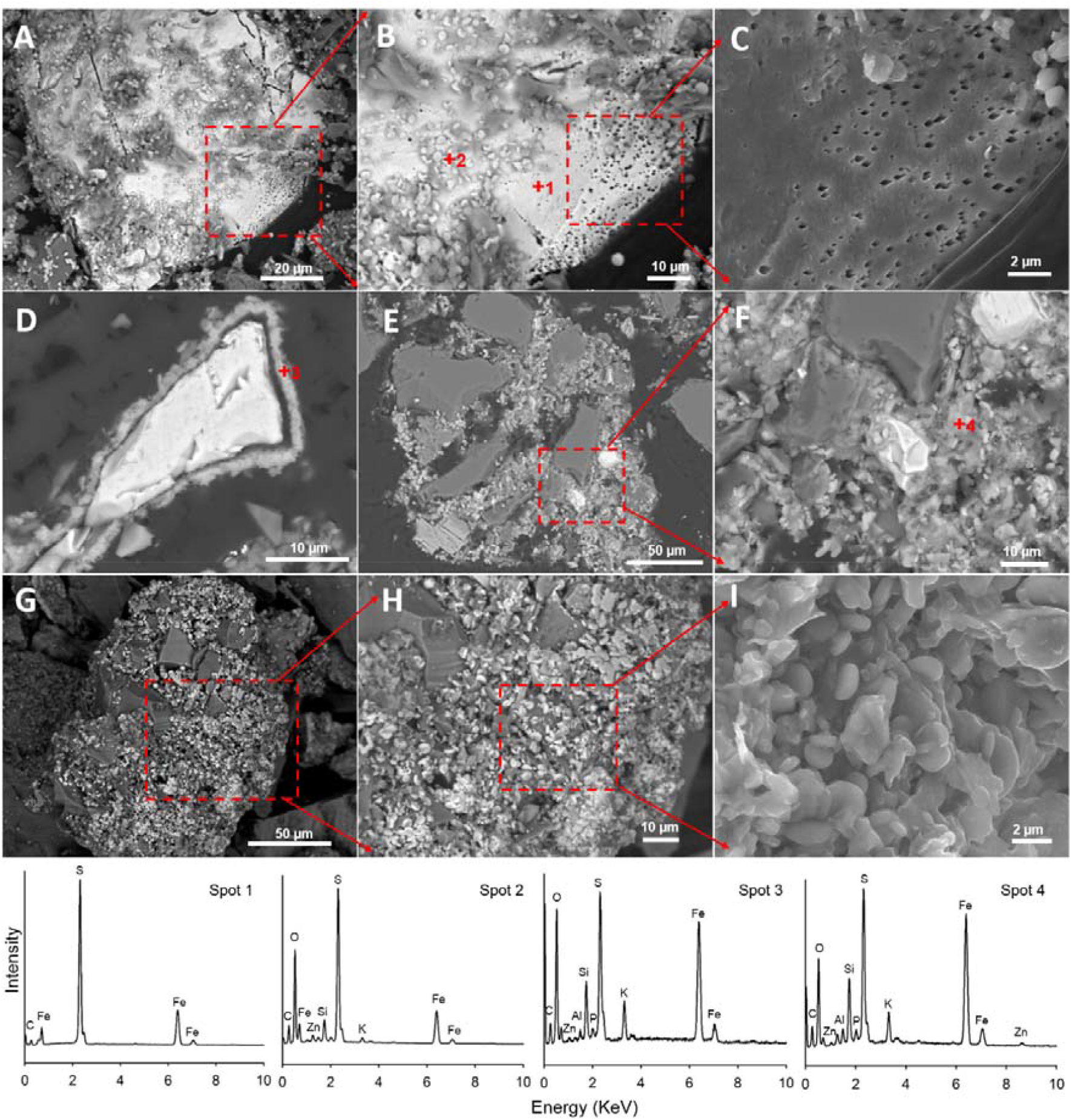
BSE-SEM-EDS showing morphology and elemental composition of Fe-bearing minerals in the TB. Micrographs “A”, “B”, “C” are unpolished tailings showing pyrite dissolution at different magnification levels (within red rectangle areas). Micrographs “D”, “E”, “F” are polished tailings showing jarosite passivation and cementation. Micrographs “G”, “H”, “I” are unpolished tailings showing a single microaggrecate and the microstucture of jarosite cements. Note: “TB” is the treatment labelled in **Fig 1**.

### 3.5 Transformation of Zn-bearing minerals

The surface morphology of Zn-sulfide grains was modified significantly to have formed densely distributed pits in the TB treatment, but not in the TC (**Fig 4**). The bio-weathering of Zn sulfides in the TB resulted in about 10 times more water-soluble Zn (the first fraction) than that of the TC, based on Zn sequential extraction (**Fig. S9**). In contrast, the exchangeable Zn pool (second fraction) was substantially lower in the TB than the TC and T0. In the strongly reducible phases (fifth fraction, bond to crystalline Fe), Zn concentration in TB was ca. 200 mg kg^−1^, while only a few Zn (ca. 5 mg kg^−1^) was detected in the TC and T0. Accordingly, the proportion of oxidizable phase (sixth fraction) in the TB was diminished by two times compared to the TC and by five times compared to the T0.

**Fig. 4.**
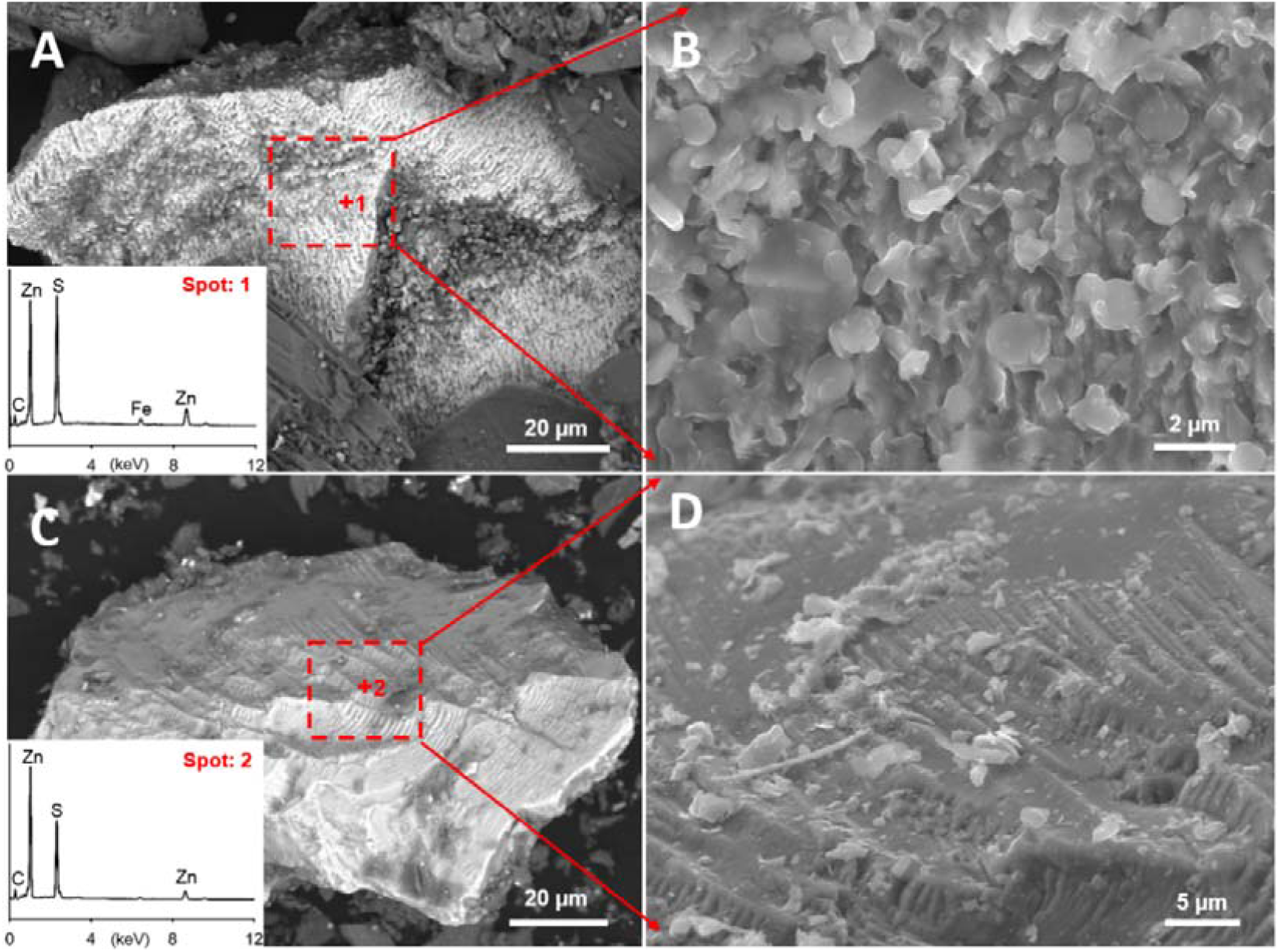
SEM micrographs highlighted that the Zn sulfide grain was being dissoved within the TB system, while the surface mophology remains unaltered in the TC. A: BSE micrograph of the Zn sulfide grain deposited on the TB; B: SE micrograph of the magnification box of “A”; C: BSE micrograph of the Zn sulfide grain deposited on the TC; D: SE micrograph of the magnification box of “C”. Note: “TB” and “TC”” are treatments labelled in **Fig 1**.

### 3.6 Zinc distribution and speciation revealed by synchrotron XFM/XANES imaging

Among the heavy metals in the tailings, the distribution and speciation of Zn (due to its abundance) were particularly investigated for verifying the concept of heavy metal encapsulation in mineral cements of the neoformed hardpans. The XFM maps suggested that Zn distribution appeared to be heterogeneous in the TB and TC, in terms of elemental intensity (**Fig. 5A**) and association (**Fig. 5B**). In the tri-colored XFM maps of Fe (red), Zn (blue) and Mn (green), the strong Zn-Fe association (purple) present in the TB demonstrated that the labile Zn might have been adsorbed onto the secondary Fe (oxyhydr)oxides (*e*.*g*., ferrihydrite or jarosite, as detected by Fe K edge XAFS in **Fig. 2**) during the bioweathering process (**Fig. 5B**). In contrast, Zn-Fe association was not evident in the TC, as little overlapped colors were displayed on the tri-colored XFM map (**Fig. S10**).

**Fig. 5.**
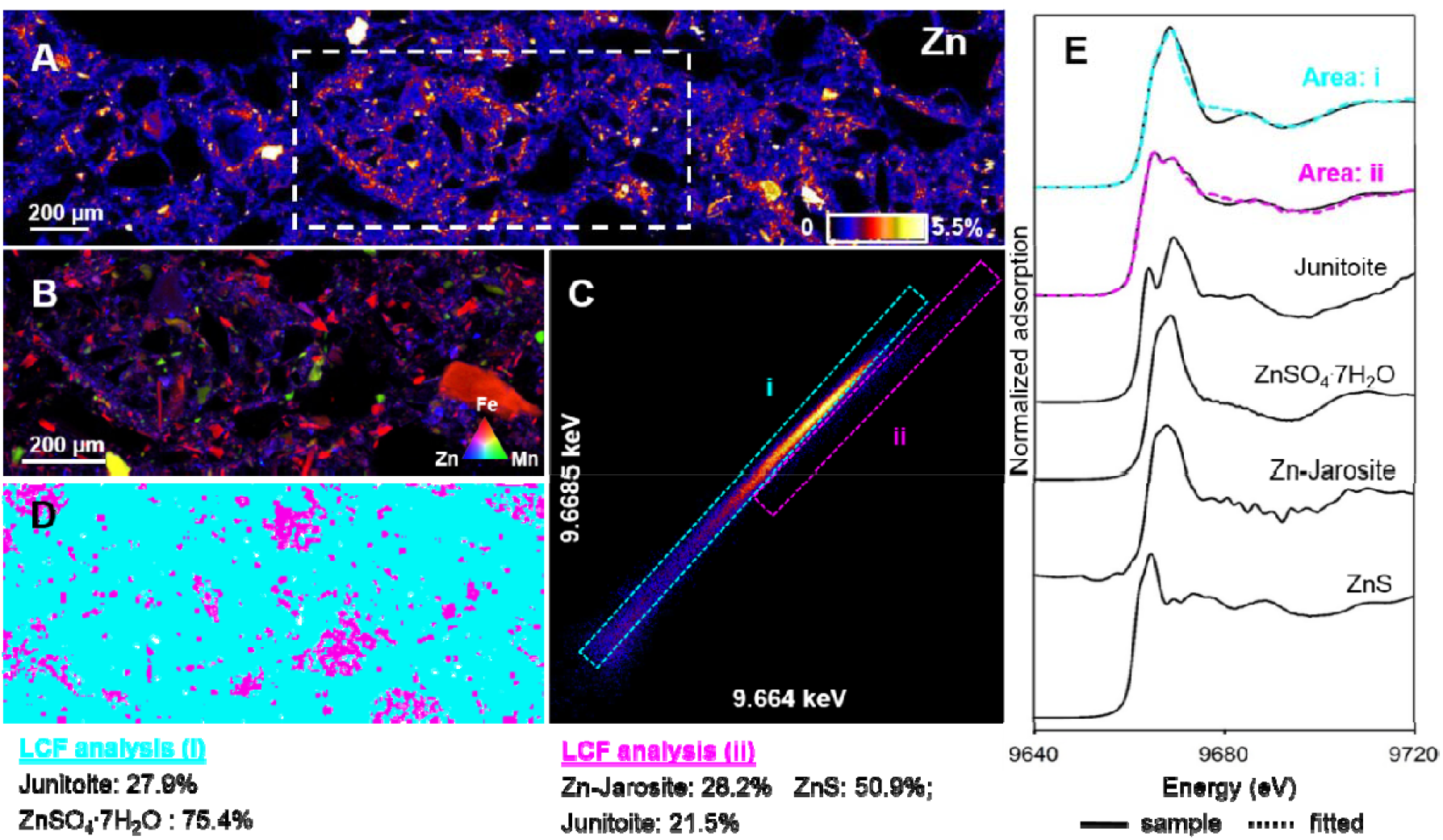
High-resolution map from XFM showing the distribution of Zn within the TB, with fluorescence-XANES imaging performed for the area indicated by the white rectangle (**A**). A tri-coloured XFM map of Fe (red), Mn (green), and Zn (blue) for the area indicated by the white rectangle in (**A**) examined by fluorescence-XANES imaging (**B**). An energy association plot from the fluorescence-XANES imaging, showing the relationship between two energies (**C**). The distribution of the two pixel populations was identified (aqua and purple) based on the dashed rectangle areas (i and ii) of the energy association plot (**D**). The XANES spectrum extracted from the pixel population shown in (**D**), plus XANES spectra for four standard compounds (**E**). Note: the sum of components was not 100% in the LCF-XANES. The r-factor, chi-square and reduced chi-square were 0.0165176, 0.63234, and 0.0076186 for Area i; 0.0031517, 0.06925, and 0.0008445 for Area ii, respectively. Note: “TB” is the treatment labelled in **Fig 1**.

The fluorescence-XANES imaging was used for the analysis of Zn speciation within the TB (white rectangle, **Fig. 5A**). Using the ‘energy association’ module in GeoPIXE, two pixel populations were identified which differed in their speciation (see dashed aqua rectangle, area ‘i’, and purple rectangle, area ‘ii’ **Fig. 5B**). It was found that the aqua pixels (area ‘i’) corresponded to the area where massive Zn distributed in the tailings, while the purple pixels corresponded to the mineral particles showing Zn-Fe associations (**Fig. 5D**). The Zn fluorescence-XANES spectra revealed distinct peak positions between the two pixel populations (**Fig. 5E**). In detail, the white line peak of aqua pixels occurred at 9668.4 eV, while the white line peak of purple pixels occurred at 9665.1 eV. Using LCF-XANES, it was predicted that the aqua area (i) was dominated by ZnSO_4_ .7H_2_O (75.4%), while Junitoite accounted for 27.9%. In contrast, the purple area (ii) was composed of ZnS (50.9%), Zn-jarosite (28.8%) and Junitoite (21.5%). In the TC, the fluorescence-XANES imaging examination of an area showed two pixel populations as well (**Fig. S10**). The LCF-XANES estimated that the aqua area (i) was dominated by ZnSO_4_ 7H_2_O (63.0%) with ZnS accounting for 38.4%. In contrast, in the purple area (ii), the Zn forms consisted of ZnS (88.7%) and ZnO (9.8%).

## 4 Discussion

As far as we have known in literature, the present study has been the first to have demonstrated the concept of bio-engineering sulfidic tailings into neoformed hardpan, by inoculating Fe/S-oxidizing bacterial consortium to accelerate the weathering of sulfides and formation of cementitious mineral gels. Synchrotron-based XAFS analysis revealed that the microbial process and associated mineral transformation in the inoculated tailings led to the formation of critical precursor mineral gels, *i*.*e*., jarosite-like minerals, which cemented the tailing minerals into integrated physical structure as the onset of hardpan formation. Meanwhile, synchrotron-based XFM analysis revealed that labile Zn liberated in the weathering were encapsulated in the jarosite-like minerals, rendering *in situ* immobilization of heavy metals in the tailings. This is consistent with our findings in field hardpan capping sulfidic Cu-Pb-Zn tailings, which naturally formed under semi-arid climatic conditions^7^.

### 4.1 Formation of secondary Fe-bearing cements accelerated by bio-weathering processes

The various microspectropic analysis revealed that large amounts of secondary Fe-bearing minerals were formed during the bio-weathering processes. The Fe oxidizing bacteria (*A. ferrooxidans*) could gain energy via Fe(II) oxidation to Fe(III) under acidic pH conditions^32, 33^, which was evidenced by the peak shifting to 7113.5 eV in pre-edge of Fe K edge XANES in the TB (**Fig. 2A**). Additionally, it is also possible that *Acidithiobacillus spp*. created a localized acidic micro-environment on biotite surfaces, thereby enhancing biotite dissolution, since biotite dissolution rate varies by up to 1.5 orders of magnitude at a given acidic pH condition^34, 35^. The weathering of biotite may release K^+^ from the interlayer of crystal^36^, which may have co-precipitated with ferric iron and sulfate to form jarosite-like minerals. Jarosite formation is regulated by microbial activities when suitable cations such as K^+^, NH_4_ ^+^, or Na^+^ are available at the pH range of 1.5-3.0 (**Eq. 1**)^37-39^ which is consistent with the observed pore-water pH conditions in the present experiment (**Fig. S4**).

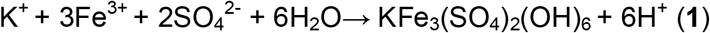

Large amounts of jarosite-like minerals may have been formed after two months of Fe/S-oxidizers colonization, as indicated by the rapid decrease of leachate K concentrations in the TB, compared to that of TC (**Fig. S6**). This supported the hypothesis that when *Acidithiobacillus spp*. oxidized ferrous iron in the presence of biotite, the dissolution and structural alteration of this trioctahedral mica released interlayer K^+^ which was concurrently incorporated as a structural cation in jarosite. The Si incorporation into the jarosite may have resulted from the co-precipitation of dissolved Si with Fe^3+^ and SO_4_^2-^. The dissolved Si should be originated from the weathering of biotite or other Si-rich minerals, as the leachate Si concentration in the TB was approximately 5 times higher than that of TC (**Fig. S6**). Structural Fe(II) in the biotite-like minerals is sensitive to redox reactions mediated by microorganisms^40^. Under the bio-oxidative conditions, the Fe^2+^ located within the biotite-like minerals can be oxidized to Fe (III) (oxyhydr)oxides^41^, leading to more ferrihydrite-like minerals formed in the TB than that in the TC. Jarosite formation requires reducing and acidic conditions, while the evaporative environments in the current condition favours jarosite preservation (*i*.*e*. cements accumulation)^38, 42^. As a result, the resultant formation of jarosite-like cements and depletion of pyrite and biotite-like minerals are primary precursors of hardpan formation, which can be accelerated by microbe driven bio-weathering processes. The accumulation of ferrihydrite and jarosite was found in the hardpan layers of sulfidic tailings, acting as cementation materials^5, 43^.The metastable jarosites may be further transformed into goethite or hematite^44, 45^. However, these reactions are more likely to occur under neutral pH rather than acidic pH^45^.

### 4.2 Zinc dissolution and mineral encapsulation within mineral cements

Zinc sulfide is acid soluble and therefore commonly depleted under acidic conditions within weathered tailings^46^. As a result, the Fe/S-oxidizing bacterial consortium induced Zn dissolution, as evidenced by the rapid release of labile Zn in the leachates (**Fig. S6**) and water-soluble Zn fraction in the TB (**Fig. S9**). The fluorescence-XANES imaging also predicted that junitoite content was lower in the TB than that of the TC, suggesting that the microbial consortium may have dissolved the Zn-Si-bearing minerals. Two possible (direct/indirect) microbial pathways may have stimulated the dissolution of Zn sulfide. In the direct pathway, *Acidithiobacillus spp*. may break up the crystalline structure through sulfur extraction or electrochemical dissolution^47, 48^, resulting in surface erosion (or pits) on the Zn sulfide grains (see the SEM images **Fig. 4**). In the indirect pathway, the microbes may firstly oxidize Fe^2+^ in bulk sulfide minerals (e.g., pyrite) and/or biotite-like minerals (**Eq. 2**) into Fe^3+^, which acts as an energy source in for the oxidizers to continue Zn sulfide oxidation^49^ (**Eq. 3**).

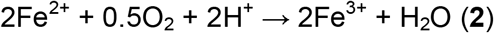

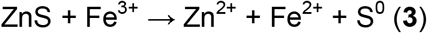

Secondary Fe (oxyhydr)oxides produced from the weathering of the sulfidic Pb-Zn tailings attenuated heavy metals across the hardpan-based tailing landscapes^50, 51^. In the present study, the SEM-EDS examination of cementation structure (**Fig. 2**) and bulk Zn sequential extraction of strongly reducible phases (fifth fraction, bond to crystalline Fe) (**Fig. 5**) highlighted the capacity of Zn attenuation by the newly formed secondary Fe (oxyhydr)oxides in the TB. The evaporative processes may facilitate heterogeneous Zn distribution in the vertical profile, with relatively more concentrated distribution in the cemented layers than un-cemented layers^7^. Using fluorescence-XANES imaging, it was found that the labile Zn in the TB was encapsulated and immobilized within mineral cements in the form of Zn-jarosite-like minerals. The Zn sulfide dissolved during the weathering processes at pH 2-3 may eventually form Zn mineral complexes with Fe (oxyhydr)oxides and jarosites in a sulfidic tailing^52^. However, no Zn-ferrihydrite-like minerals were detected in the TB even though ferrihydrite usually indicated a higher heavy metal sorption capacity than jarosite due to its large surface area^53^. Our previous research demonstrated that the neutral pH condition in the hardpan formed at the surface sulfidic Cu-Pb-Zn tailings could restrict Zn co-dissolution with ferrihydrite and re-distribution to other Fe-bearing minerals^7^. In contrast, the labile Zn initially adsorbed at the ferrihydrite surfaces may be co-dissolved with ferrihydrite at acidic pH induced by bacterial inoculation^54^, which may be selectively adsorbed on jarosite surfaces. The formation of crystalline (or poorly-crystalline) jarosite-like mineral gels would result in irreversible metal sorption or a permanent sink for sorbed metals in the tailings^55, 56^. The inert geochemical reactivity and encapsulation of heavy metals mobilized by jarosite-like mineral gels add the advantage to the proposed hardpan capping method for minimizing long-term pollution risks of the hardpan layers.

### 4.3 Environmental implications

Building on the wealth of knowledge about the characterization of acidic/neutral and metallic drainages in sulfidic tailings, the present study has advocated a proactive approach of *in situ* microbial weathering and hardpan capping method to meet environmental challenges of billions of tones of sulfidic and metallic tailings stored across geographical locations worldwide. Through microbial regulation of the underlying processes in the formation of secondary jarosite-like mineral gels, this concept-proven bio-engineering process may be scaled up in further field trials to develop this novel cover method of *in situ* formation of massive hardpan caps. These hardpan caps can form duplex-soil systems together with reconstructed root zones and sustain native plant communities across sulfidic tailing landscapes for closure outcomes.

## Supporting information

Information regarding the bulk geochemical properties, methods for bacterial inoculum preparations, XAFS reference standards preparation, SEM-EDS imaging, pH, EC, and heavy metal concentration in the leachate, and the data for abiotic control tailings are available in **SI**.

## Acknowledgements

This research was undertaken on the XAS beamline 01C1 at National Synchrotron Radiation Research Centre, Taiwan and XFM beamline at the Australian Synchrotron (AS182/XFM/13331), part of ANSTO. The authors also acknowledge the facilities, and the scientific and technical assistance, of the Australian Microscopy & Microanalysis Research Facility at the Centre for Microscopy and Microanalysis, The University of Queensland (UQ). The study was supported by UQ research higher degree grant (9342909-01-205-21).

