## Supporting information for "Microbial activated mineral weathering and cementation as precursors to hardpan formation and heavy metal encapsulation in sulfidic tailings"

Number of Pages: 20

Number of Figures: 15

Number of Tables: 2

**Bacteria enrichment methods**

*Acidithiobacillus ferrooxidans* (DSM 14882) and *Acidithiobacillus thiooxidans* (ATCC 19377) cultures were incubated in basal media supplemented with ferrous iron and elemental sulphur, respectively, at 30°C in a rotary shaker at 120 rpm. *A. ferrooxidans* was cultured in 9K nutrient medium containing $(NH_4)_2SO_4$ (0.4 g), $K_2HPO_4$ (0.1 g), $MgSO_4 \cdot 7H_2O$ (0.4 g), $FeSO_4 \cdot 7H_2O$ (20 g) in one litre of deionised water at pH 2.3 (Silverman and Lundgren, 1959). *A. thiooxidans* was cultured in $(NH_4)_2SO_4$ (0.3 g), $K_2HPO_4$ (0.1 g), $MgSO_4 \cdot 7H_2O$ (0.4 g), $CaCl_2 \cdot 2H_2O$ (0.33 g) and S (ca. 1 g) in one litre of deionised water at pH 2.3 (Starkey, 1925).

**Fe K-edge XAFS Analysis Methods**

Fe K-edge (7,112 eV) XAFS spectra were collected on beamline 01C1 at the national synchrotron radiation research center (NSRRC) in Taiwan. The storage ring was operated at 1.5 GeV, and the ring current was in the 360 mA. We selected the following Fe standards (1) magnetite, (2) hematite, (3) goethite, (4) ferrihydrite, (5) siderite, (6) Fe(III) phosphate, (7) biotite, (8) olivine, (9) pyrite, (10) illite, (11) jarosite, (12) vermiculite, (13) Fe(II) gluconate hydrate, (14) Fe(II) glutathione, (15) Fe(III) oxalate, (16) Fe(II) humic acid (17) Fe(II) histidine (18) Fe(III) histidine, (19) Fe(II) oxide, (20) Fe (II)-sulfate. Magnetite, hematite, goethite, pyrite, siderite, vermiculite, Fe(III) phosphate, Fe(II) oxide, Fe (II)-sulfate and Iron(II) gluconate hydrate were bought from Sigma Aldrich; Olivine, illite and biotite were purchased from Ward's science (https://www.wardsci.com); jarosite was purchased from Kremer Pigmente (https://www.kremer-pigmente.com/en); Ferrihydrite (2-line) was synthesized according to the method developed by Schwertmann (2008). Briefly, 40 g $Fe(NO_3) \cdot 9H_2O$ was dissolved in 500 ml MilliQ water and 1.0 M KOH was added to adjust the pH to 7-8. The products were centrifuged, dialysed and freeze-dried for use. Fe-organic matter complexes were prepared through mixing 0.1 M Fe(II)/Fe(III) solutions with organic matters (*i.e.,* oxalate acid, humic acid, histidine, glutathione at a ratio of 1:5 mole (Guénet et al., 2017).

An energy range of -200 to 800 eV relative to the Fe K-edge absorption was used to acquire spectra of tailing/hardpan and solid Fe reference compounds in transmission mode under ambient conditions. Fe K-edge XAS spectra of Fe organic matter complexes were collected at fluorescence mode. Data were collected using the following energy ranges: -200 to - 20 eV (10 eV steps with 2s per interval); -20 - 30 eV (0.35 eV steps with 2s per interval); 30 - 800 eV (4 eV steps with 4s per interval).

The energy scale was calibrated using a Fe foil as an internal standard (calibration energy is 7112.0 eV). The foil was placed between the I1 and I2 ion chambers and collected simultaneously with each sample spectra. The XAS data were normalized (baseline and background corrections) using ATHENA from DEMETER (or IFEFFIT) software package (CARS, University of Chicago) (Ravel and Newville, 2005).

**Table S1:** Background geochemistry properties of the tailings using for this study (Individual values were mean ± standard deviation, triplicates).

| | pH[a] | EC[b] |
|---|---|---|
| | 6.9±0.3 | 0.6±0.1 |
| | (Pseudo)-total[c] (mg/kg) | Water soluble (mg/kg) |
| Al | 11471±382 | 2±0.2 |
| As | 983±21 | U.D[d] |
| Ca | 25510±1012 | 672±72 |
| Cd | 9±1 | U.D |
| Co | 5±2 | 4±1 |
| Fe | 77990±3212 | U.D |
| K | 1868±14 | 71±6 |
| Mg | 9637±89 | 21±3 |
| Mn | 3333±56 | 1±0.2 |
| Na | 537±22 | 227±13 |
| Pb | 4162±44 | 1±0.3 |
| S | 16158±1032 | 548±34 |
| Zn | 4263±432 | 2±0.5 |

[a] Measured with 1:5 solid/water suspension
[b] Electrical Conductively
[c] Bulk tailings were determined by an ICP-OES following *aqua regia* digestion
[d] Under detection limit of the ICP-OES

The pH and electrical conductivity (EC) in the samples were measured with 1:5 solid/water suspensions by using a bench-top pH-Conductivity meter (TPS 901-CP, Brisbane, Australia). Pseudo-total concentrations of elements concerned in the bulk samples were determined using an inductively coupled plasma optical emission spectroscopy (ICP-OES) (PerkinElmer Optima 8300, Waltham, USA) following *aqua regia* digestion. Water-extractable elements in the hardpan and tailings were measured as previously described (Dold 2003).

**Table S2**: Seven steps sequential extractions scheme used to investigate the distribution of chemical forms of metals in the tailings. (Modified from (Tessier et al., 1979; Sondag, 1981).

| Extraction step | Solvent | Target phases |
|---|---|---|
| 1 | Deionised water; 1 hour end-over-end shaking | Water-soluble: Evaporitic minerals, e.g. halite (NaCl), gypsum ($CaSO_4.2H_2O$) |
| 2 | 1 M ammonium acetate solution pH 4.5; 2 hours end-over-end shaking, at room temperature | Exchangeable: Calcite, vermiculite-type mixed-layer, adsorbed and exchangeable ions |
| 3 | 1 M sodium acetate ($CH_3COONa$) solution adjusted to pH 5 with acetic acid ($CH_3COOH$); 2 hours end-over-end shaking | Weakly acid-soluble: Carbonates |
| 4 | 0.2 M ammonium oxalate ($(NH_4)_2C_2O_4$) solution adjusted to pH 3.0 with 0.2 M oxalic acid ($H_2C_2O_4$); 1 hour end-over-end shaking in darkness | Weakly reducible: Amorphous or poorly crystalline oxyhydroxides of $Fe^{3+}$ and $Mn^{3+}$ (e.g. ferrihydrite) |
| 5 | 0.2 M ammonium oxalate solution adjusted to pH 3.0 with 0.2 M oxalic acid; 2 hours in 80°C water bath with vortex mixing every 30 minutes | Strongly reducible: Crystalline $Fe^{3+}$ oxides (e.g. goethite, hematite), jarosite, Al oxyhydroxides |
| 6a | Potassium chlorate ($KClO_3$), 10 ml concentrated HCl and 25ml deionized water; 45 minutes standing in fume hood, or until reaction no longer visible | } Sulfides (e.g. ZnS, PbS, $FeS_2$) |
| 6b | 25ml 4M nitric acid ($HNO_3$); 40 minutes in boiling water bath. | |
| 7 | Aqua regia digestion ($HCl+HNO_3$) | Residual: Silicates and non-silicate phases occluded by silicates. |

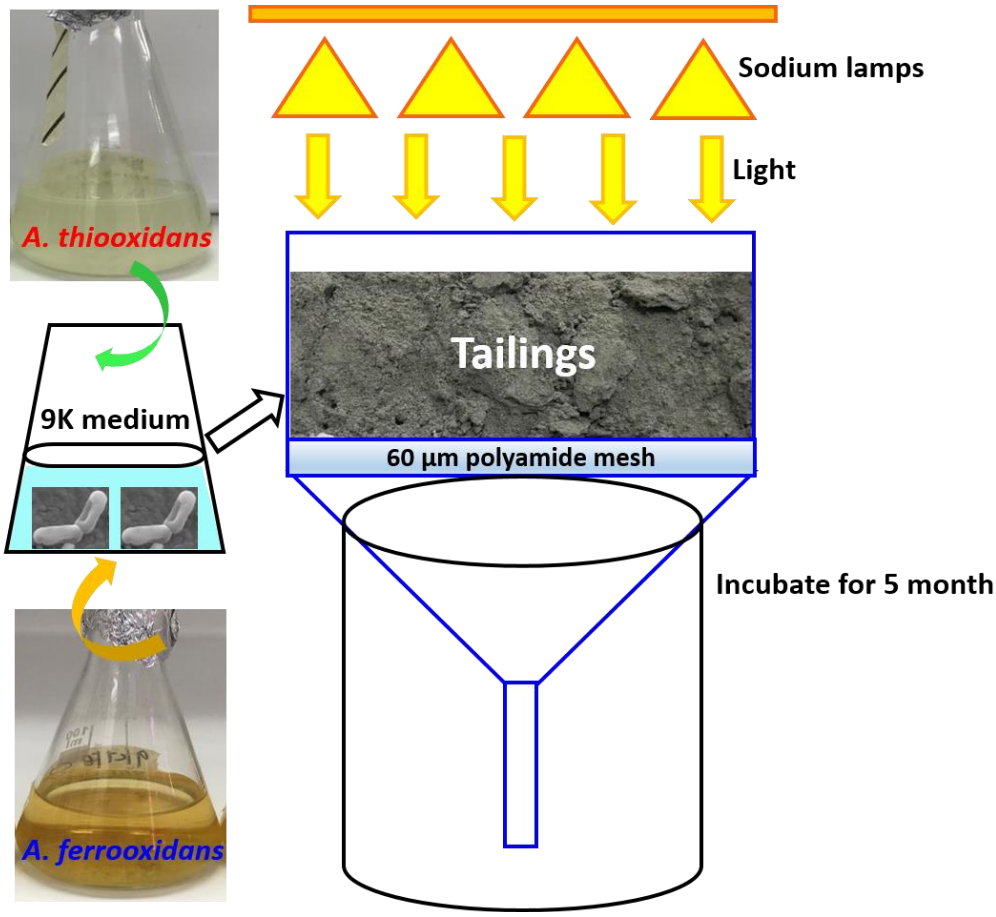


**Fig. S1:** The microbial inoculum preparation and funnel column set-up in this study.

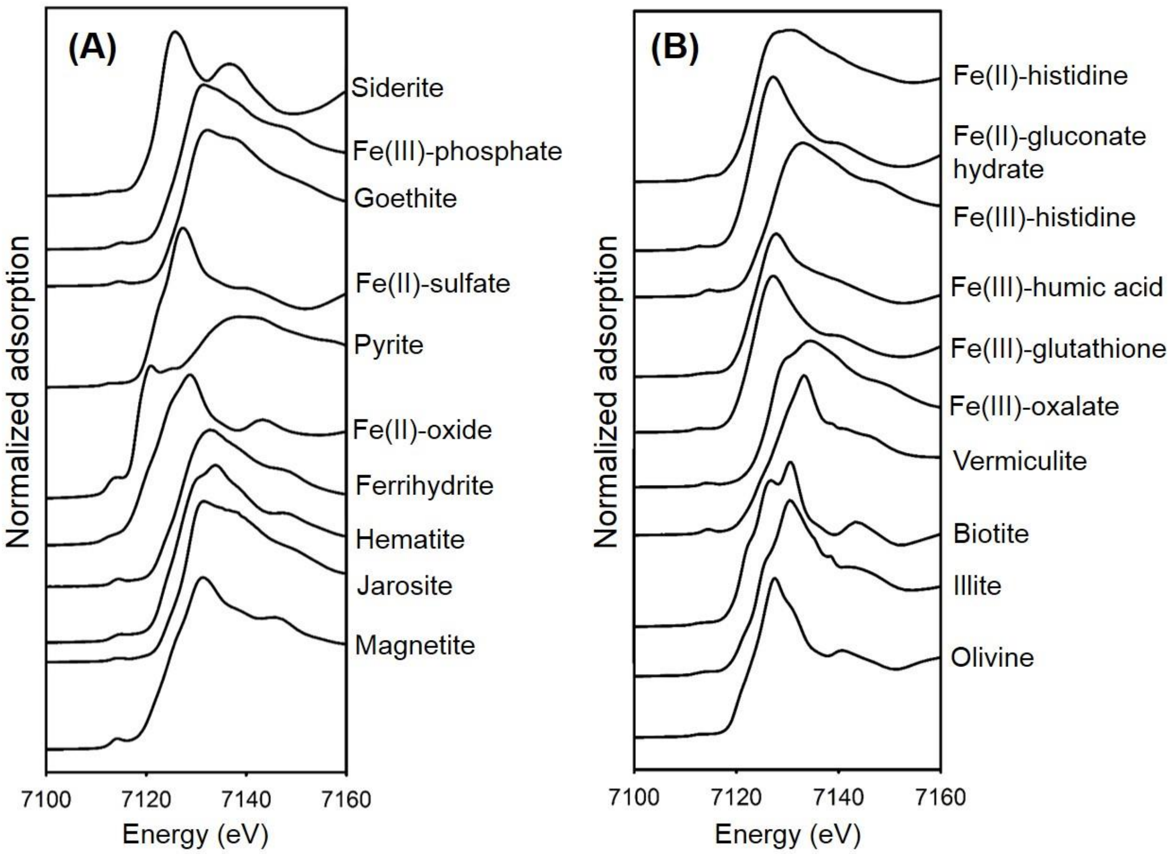


91

92 **Fig. S2** XANES spectra of twenty Fe reference standards analysed in this study.

93

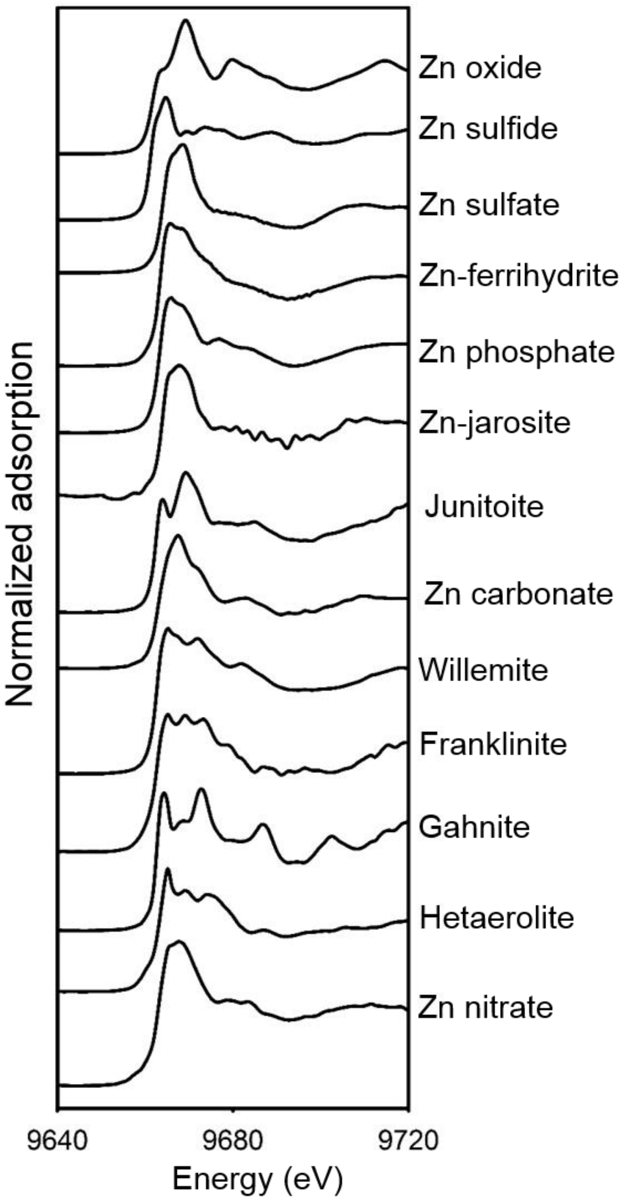


94 **Fig. S3** XANES spectra of Zn standard compounds analysed in this study.

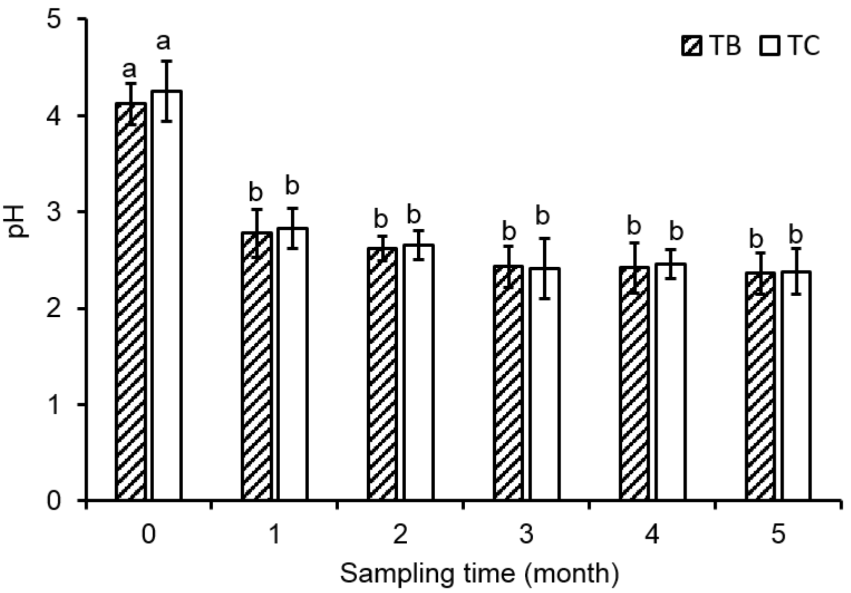


**Fig. S4** pH in the leachate during the 5-month incubation period. Different letters indicate significant differences of corresponding means among the treatments (one-way ANOVA followed by LSD test, $p < 0.05$). Note: "TB" represents tailings inoculated by the bacterial consortium (containing both *A. thiooxidans* and *A. ferrooxidans*); "TC" represents the tailings receiving the same medium without bacterial inoculation as the control.

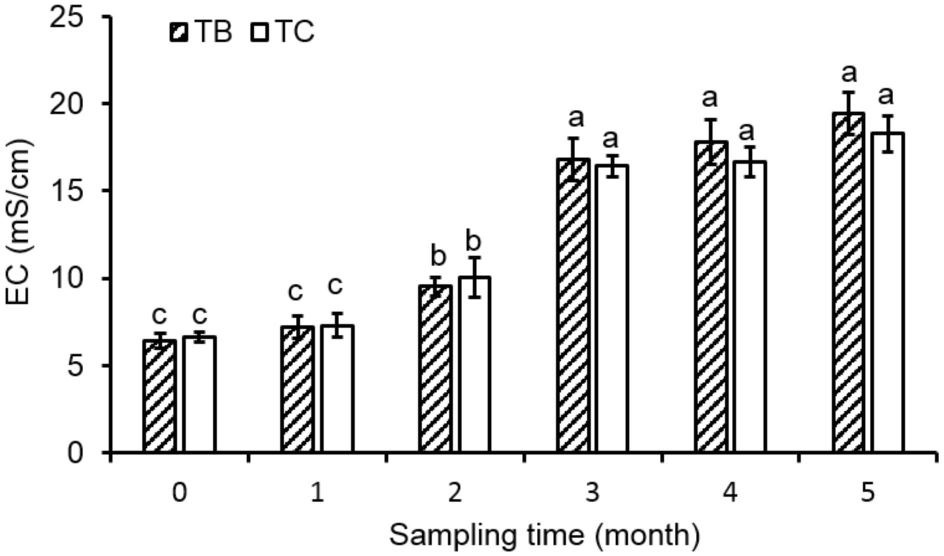


**Fig. S5** EC in the leachate during the 5-month incubation period. Different letters indicate significant differences of corresponding means among the treatments (one-way ANOVA followed by LSD test, $p < 0.05$). Note: "TB" represents tailings inoculated by the bacterial consortium (containing both *A. thiooxidans* and *A. ferrooxidans*); "TC" represents the tailings receiving the same medium without bacterial inoculation as the control.

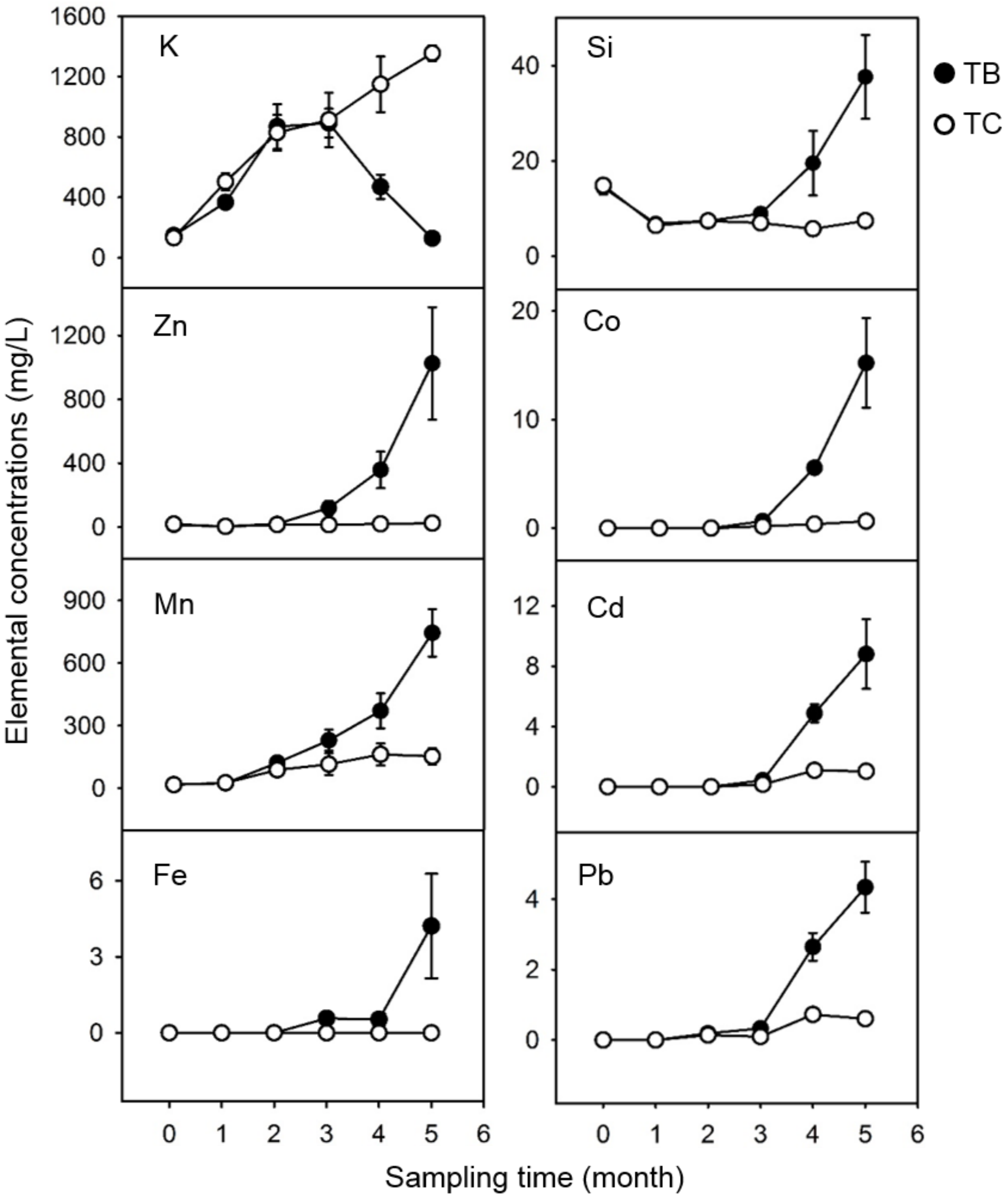


**Fig. S6** Elemental concentrations in the leachate during the 5-month incubation period. Note: “TB” represents tailings inoculated by the bacterial consortium (containing both *A. thiooxidans* and *A. ferrooxidans*); “TC” represents the tailings receiving the same medium without bacterial inoculation as the control.

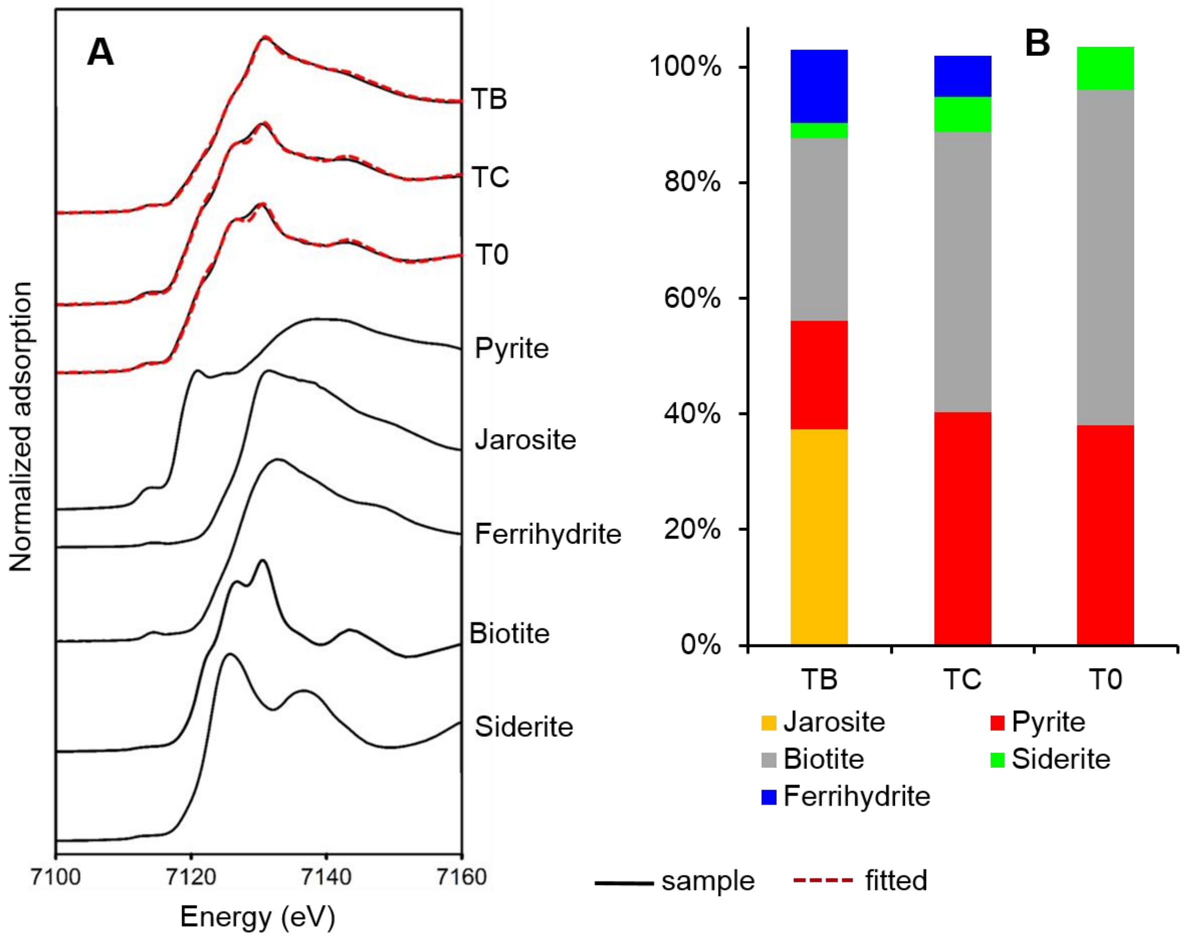


**Fig. S7** Fe K-edge XANES spectra (line) and LCF (dashed) highlight that the distinct Fe-bearing phases within the TB, TC and T0. Note: the sum of components was not 100% in the LCF-XANES. The r-factor, chi-square and reduced chi-square were 0.0003176, 0.01146, and 0.0000903 for TB; 0.006390, 0.02039, and 0.00001593 for TC; 0.0009618, 0.03265, and 0.002531 for T0, respectively. Note: "TB" represents tailings inoculated by the bacterial consortium (containing both *A. thiooxidans* and *A. ferrooxidans*); "TC" represents the tailings receiving the same medium without bacterial inoculation as the control; "T0" represents the time "0" tailings (original tailings).

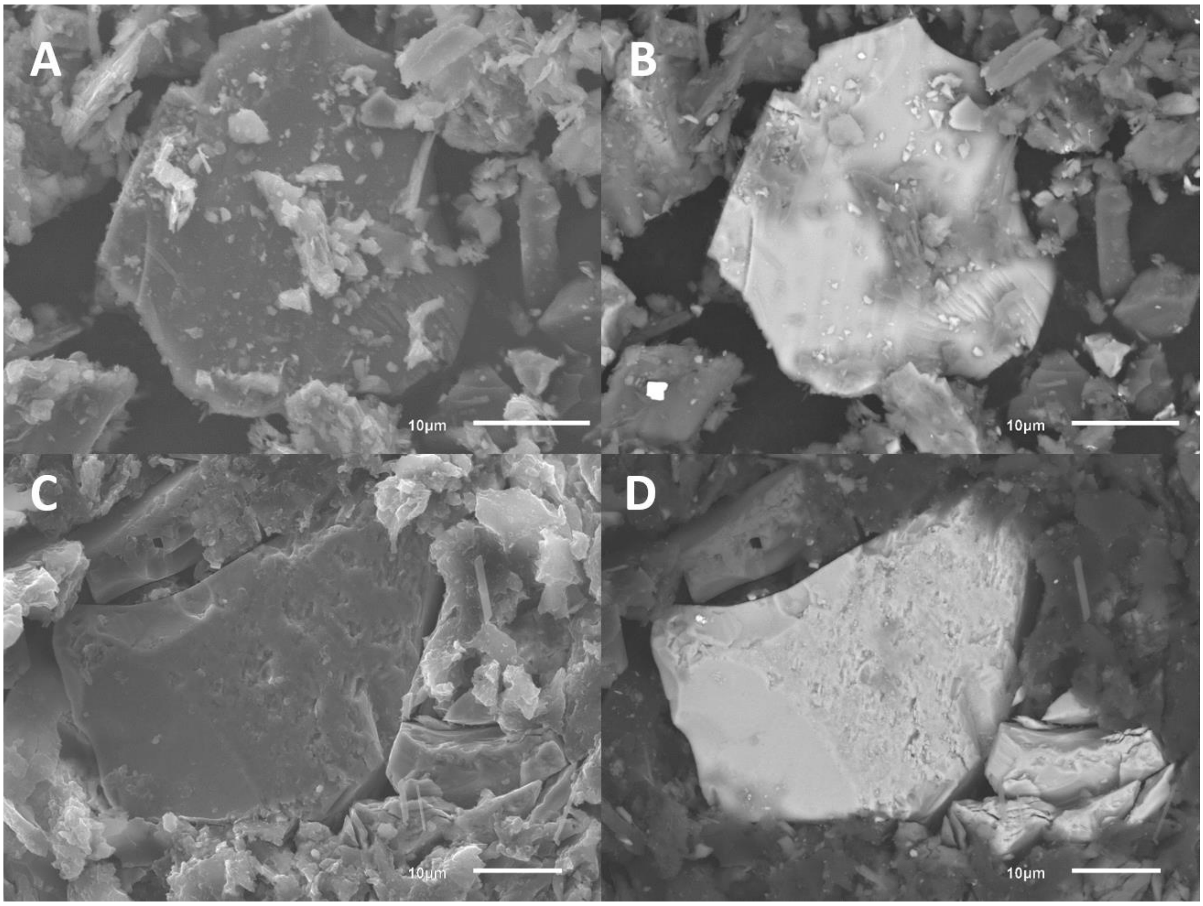


**Fig. S8** The SE micrographs (**A**) and BSE micrographs of (**B**) highlight that the surface morpology of the pyrite grain remain unchanged in the TC (unpolished). The SE micrographs (**C**) and BSE micrographs (**D**) highlight that no coating materials are deposited on the rims of the pyrite grain in the TC (polished). Note: “TB” represents tailings inoculated by the bacterial consortium (containing both *A. thiooxidans* and *A. ferrooxidans*); “TC” represents the tailings receiving the same medium without bacterial inoculation as the control.

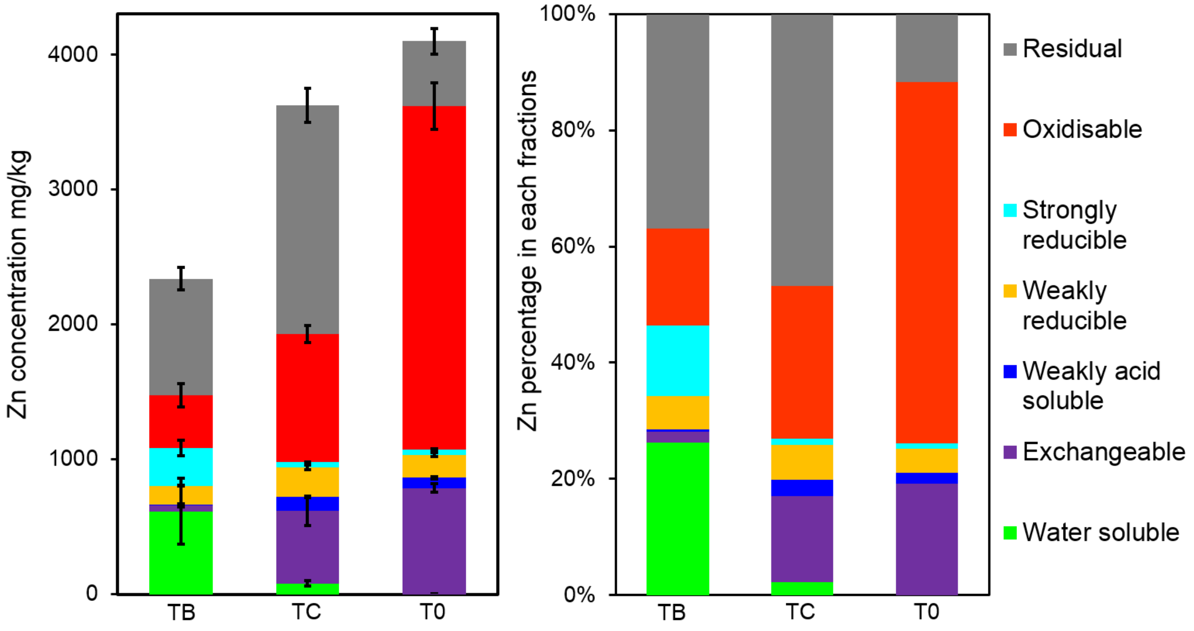


**Fig. S9** Total concentrations and percentages of Zn distributed in each fraction in the bulk tailings, as revealed by the sequential extraction method. Note: "TB" represents tailings inoculated by the bacterial consortium (containing both *A. thiooxidans* and *A. ferrooxidans*); "TC" represents the tailings receiving the same medium without bacterial inoculation as the control; "T0" represents the time "0" tailings (original tailings).

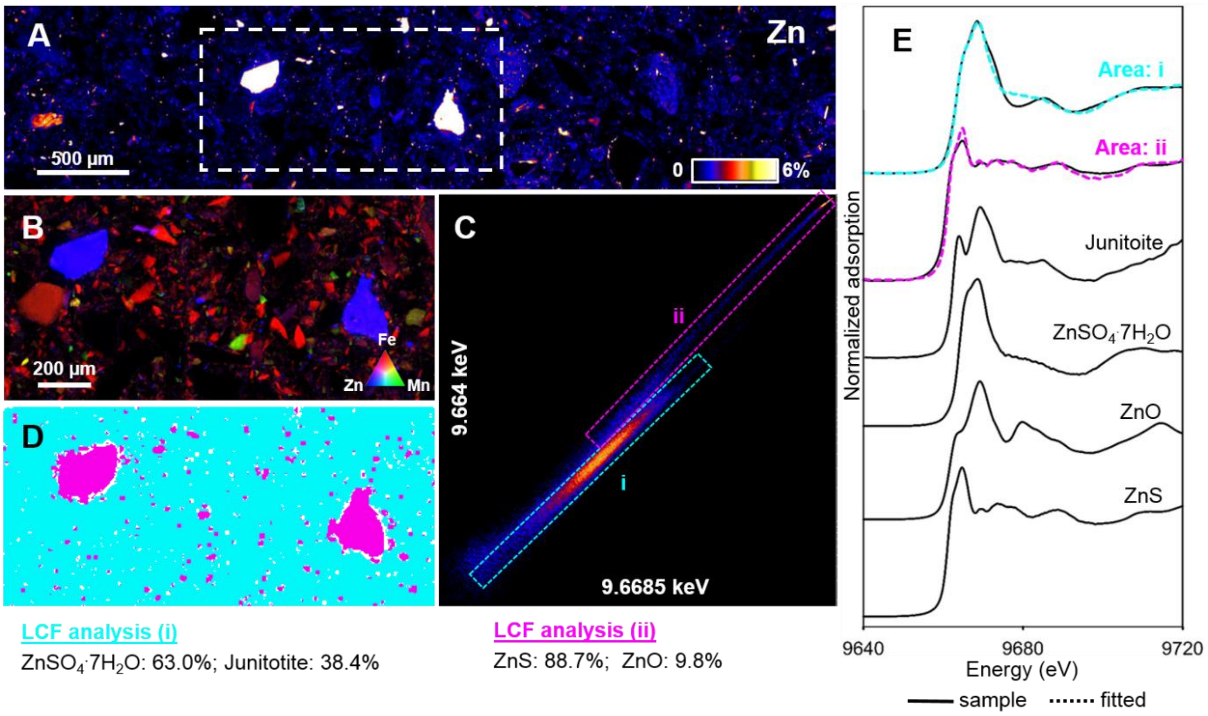


**Fig. S10** High-resolution map from XFM showing the distribution of Zn within the TC, with fluorescence-XANES imaging performed for the area indicated by the white rectangle (**A**). A tri-colored XFM map of Fe (red), Mn (green), and Zn (blue) for the area indicated by the white rectangle in (**A**) examined by fluorescence-XANES imaging (**B**). An energy association plot from the fluorescence-XANES imaging, showing the relationship between two energies (**C**). The distribution of the two pixel populations was identified (blue and purple) based on the dashed rectangle areas (i and ii) of the energy association plot (**D**). The XANES spectrum extracted from the pixel population shown in (**D**), plus XANES spectra for four standard compounds (**E**). Note: the sum of components was not 100% in the LCF-XANES. The r-factor, chi-square and reduced chi-square were 0.0116524, 0.40002, and 0.0046514 for Area i; 0.0082837, 0.12764, and 0.0015566 for Area ii, respectively.

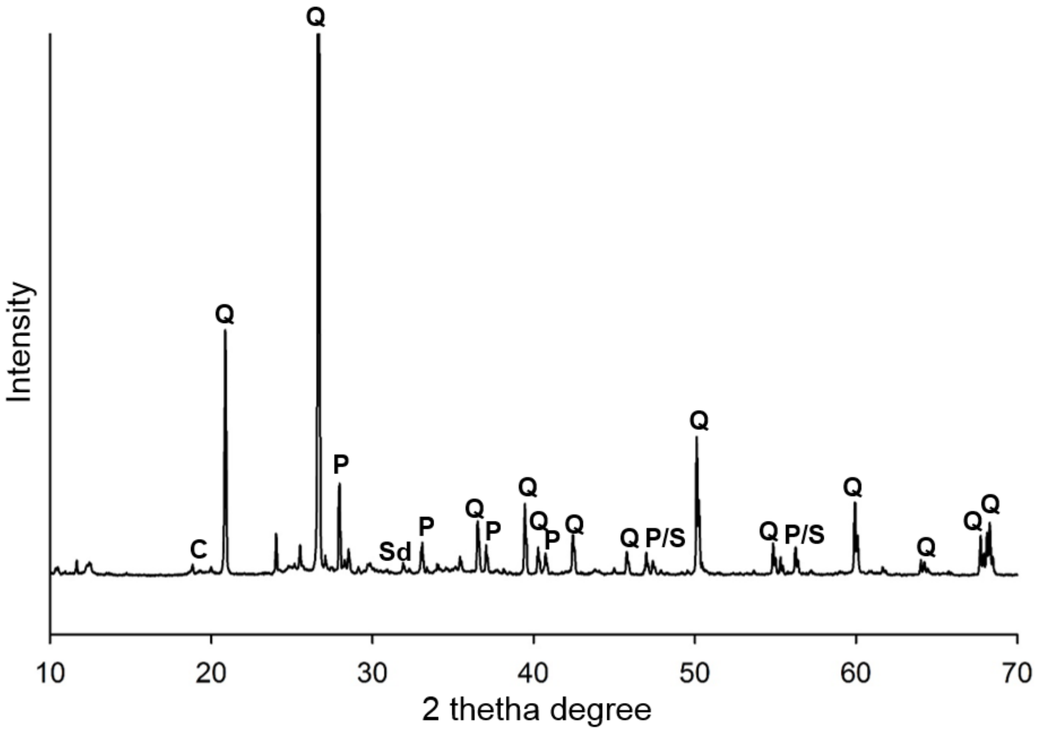


**Fig. S11** XRD spectra of the abiotic control tailings. Note: C: Calcite ($CaCO_3$); Q: Quartz ($SiO_2$); P: Pyrite ($FeS_2$); S: Sphalerite (ZnS); Sd: Siderite ($FeCO_3$).

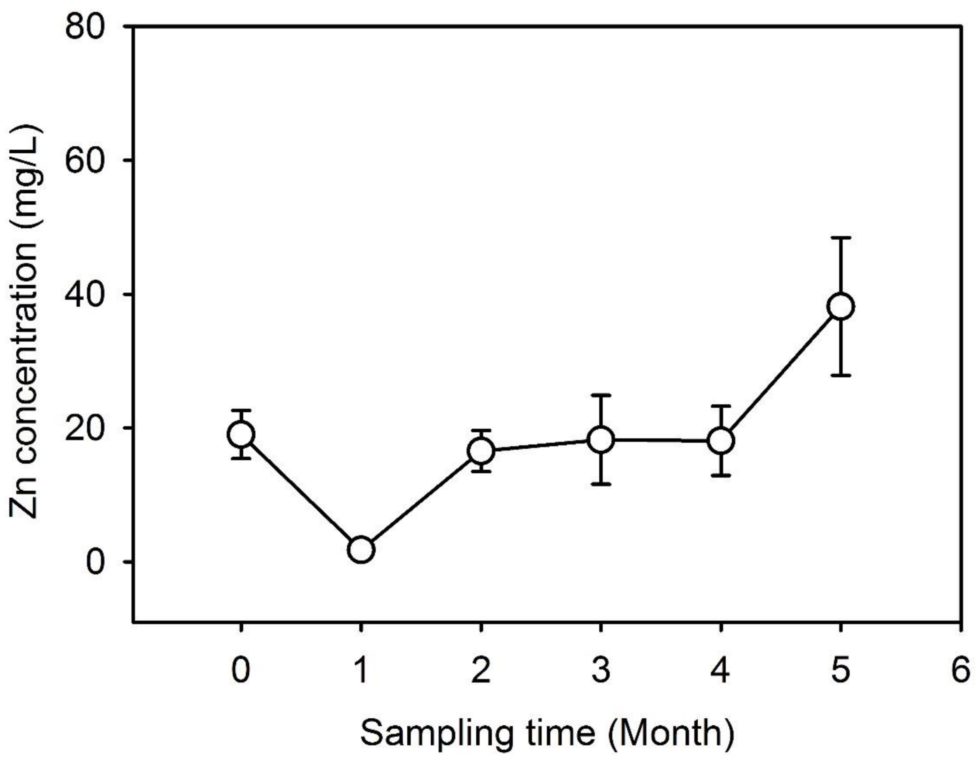


**Fig. S12** Zn concentration in the leachate from the abiotic control tailings. The Zn concentration is significantly lower than the TB, which is also corresponded to that in the TC.

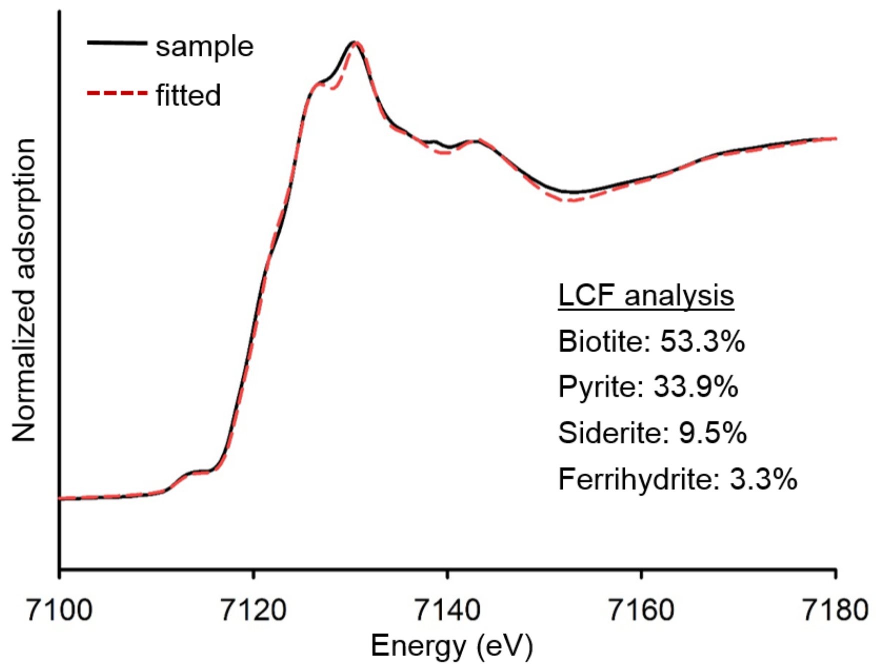


**Fig. S13** Fe K-edge XANES spectra (line) and LCF (dashed) highlight that the speciation of Fe-bearing minerals in the abiotic control is corresponded to that in the TC and T0. As for the quality control, the r-factor, chi-square, and reduced chi-square are 0.0012732, 0.02895, and 0.0002731, respectively

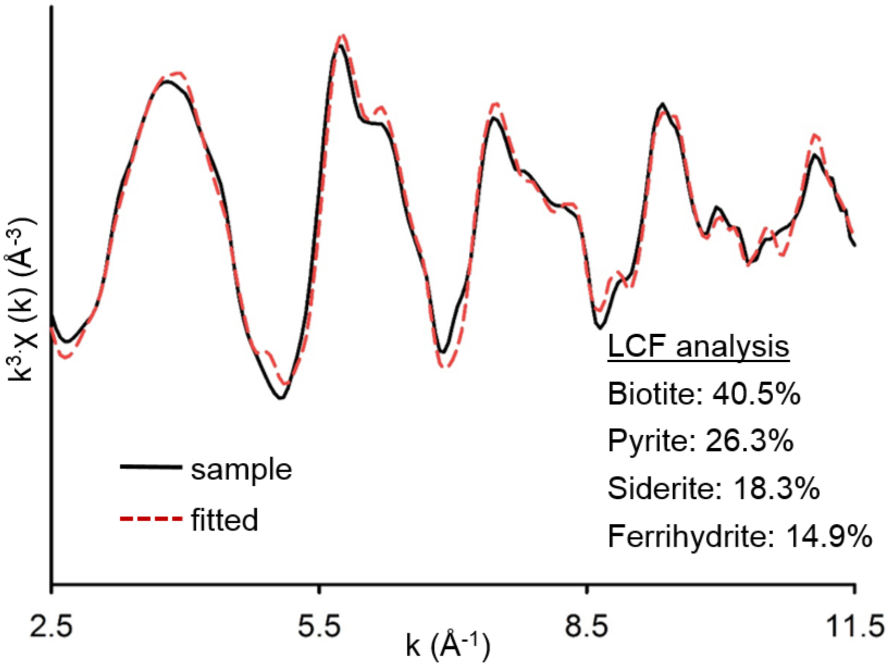


**Fig. S14** *k* space Fe K-edge EXAFS spectra highlight that the speciation of Fe-bearing minerals in the abiotic control tailings is corresponded to that in the TC and T0. As for the quality control, the r-factor, chi-square, and reduced chi-square are 0.0248459, 0.60717, and 0.0034111, respectively.

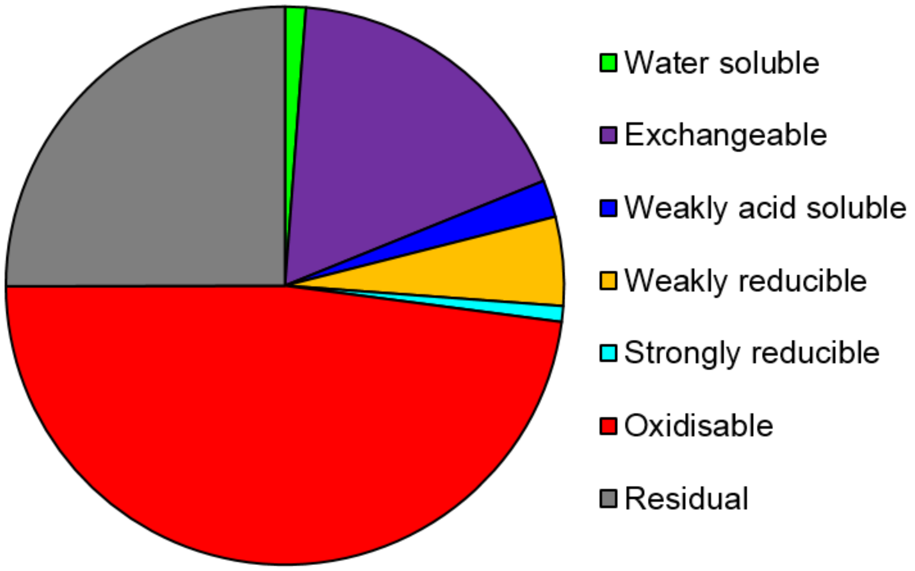


**Fig. S15:** The percentages of Zn distributed in each fraction in the abiotic control tailings, as revealed by the sequential extraction method. The results are corresponded to that of the TC and T0.
